# Multi T1-weighted contrast imaging and T1 mapping with Compressed sensing FLAWS at 3T

**DOI:** 10.1101/2021.12.18.473283

**Authors:** J. Beaumont, J. Fripp, P. Raniga, O. Acosta, J.C. Ferre, K. L. McMahon, J. Trinder, T. Kober, G. Gambarota

## Abstract

The Fluid And White matter Suppression (FLAWS) MRI sequence allows for the acquisition of multiple T1-weighted contrasts in a single sequence acquisition. However, its acquisition time is prohibitive for use in clinical practice when the k-space is linearly downsampled and reconstructed using the Generalized Autocalibrating Partially Parallel Acquisition (GRAPPA) technique. This study proposes a FLAWS sequence optimization tailored to allow for the acquisition of FLAWS images with a Cartesian phyllotaxis k-space undersampling and compressed sensing (CS) reconstruction at 3T. The CS FLAWS sequence parameters were determined using a method previously employed to optimize FLAWS imaging at 1.5T and 7T. *In-vivo* experiments show that the proposed CS FLAWS optimization allows to reduce the FLAWS sequence acquisition time from 8 *mins* to 6 *mins* without decreasing the FLAWS image quality. In addition, this study demonstrates for the first time that T1-weighted imaging with low B1 sensitivity and T1 mapping can be performed with the FLAWS sequence at 3T for both GRAPPA and CS reconstructions. The FLAWS T1 mapping was validated using *in-silico, in-vitro* and *in-vivo* experiments with comparison against the inversion recovery turbo spin echo and MP2RAGE T1 mappings. These new results suggest that the recent advances in FLAWS imaging allow to combine the MP2RAGE imaging benefits (T1-weigthed imaging with low B1 sensitivity and T1 mapping) and with the previous version of FLAWS imaging benefits (multi T1-weighted contrast imaging) in a single 6 *mins* sequence acquisition.

**Graphical abstract:** 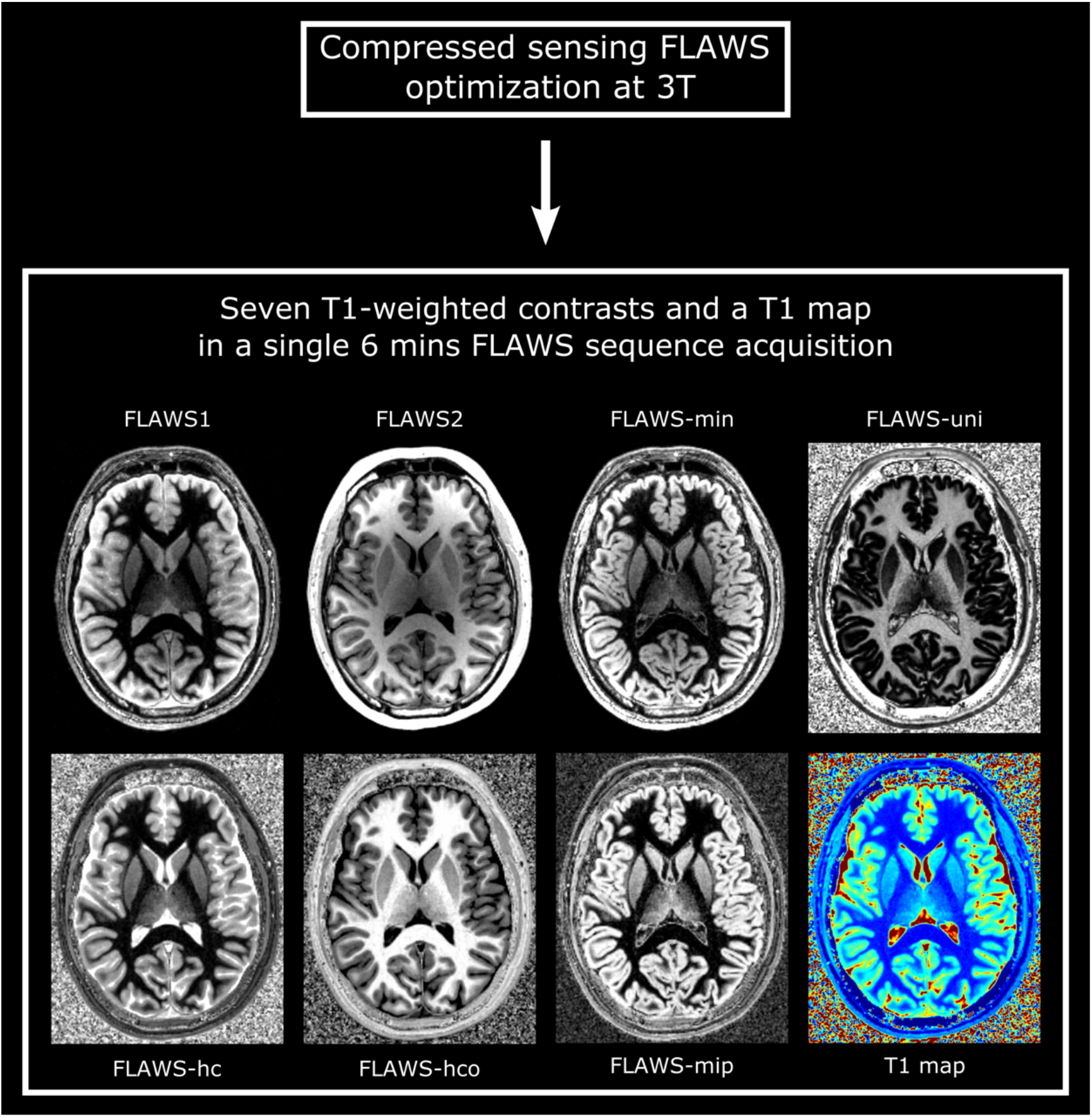

## 1. Introduction

The acquisition of magnetic resonance (MR) images from inversion recovery-based T1-weighted sequences is nowadays a gold standard to study the human brain. Specifically, the magnetization prepared rapid gradient echo (MPRAGE) sequence [1] provides a cerebrospinal fluid- (CSF) suppressed T1-weighted contrast that is routinely acquired to visualize the human brain structure. In addition, the fast gray matter acquisition T1 inversion recovery (FGATIR) sequence is used as it provides a T1-weighted contrast with white matter (WM) signal suppression that is of interest for the visualization of deep gray matter (GM) structures in applications such as deep brain stimulation (DBS) surgery planning. The CSF suppressed T1-weighted contrast provided by the magnetization prepared two rapid gradient echoes (MP2RAGE) sequence [2] usually replaces the standard MPRAGE contrast for acquisitions performed at ultra-high fields (7T and above) as it is characterized by a low sensitivity to *B*1 field inhomogeneities. This contrast, named *MP2RAGE-uni*, is obtained by combining the two naturally co-registered gradient echoes that are acquired after the inversion of the longitudinal magnetization. The MP2RAGE-uni contrast is also characterized by a low sensitivity to *M*0 and *T*2* relaxation time, thus allowing for the measurement of T1 relaxation times by using lookup tables built from the solutions of the MP2RAGE sequence Bloch equations [2].

Following its precursors, the fluid and white matter suppression (FLAWS) sequence was proposed by adapting the MP2RAGE sequence parameters in order to provide both a WM- and a CSF-suppressed contrast in a single sequence acquisition [3]. The WM-suppressed and CSF-suppressed contrasts, respectively named *FLAWS1* and *FLAWS2*, are naturally co-registered and are visually similar to the FGATIR and the MPRAGE contrasts [3]. Thanks to their co-registration properties, the *FLAWS1* and *FLAWS2* images can be combined with a voxel-wise minimum computation to provide a GM-specific contrast named *FLAWS-min* [3]. The aforementioned contrasts provided by the FLAWS sequence were shown to be of interest for multiple brain imaging applications, such as epilepsy lesion detection [4–6], deep brain structures visualization for DBS surgery planning [7,8] and brain tissue segmentation [9].

Despite an increasing interest of the brain imaging community towards the FLAWS sequence over the past few years, the use of FLAWS imaging is still limited in routine clinical practice due to its long acquisition time, which is 10 mins at 3T [3]. In a recent study conducted by Mussard et al. at 3T [10], the linear k-space undersampling combined with a GRAPPA reconstruction [11] traditionally employed in MP2RAGE imaging were replaced by a Cartesian phyllotaxis k-space undersampling [12] and compressed sensing reconstruction [13] to speed up the MP2RAGE acquisition time from 8 mins to up to 3 mins, while obtaining images of similar quality and close T1 measurements. Given the similarity between the MP2RAGE and FLAWS sequences, the method proposed by Mussard et al. to decrease the MP2RAGE sequence acquisition time may also be used to decrease the FLAWS sequence acquisition time.

Although the FLAWS sequence provides an increased clinical interest compared to the MP2RAGE sequence since it yields multiple T1-weighted contrasts, it could not replace standard MP2RAGE protocols as the *FLAWS-uni* image -the equivalent of the *MP2RAGE-uni* image for a FLAWS acquisition-does not produce a CSF-suppressed T1-weighted contrast and cannot be used to measure T1 relaxation times. In recent studies conducted at 1.5T [14,15], a new FLAWS image combination named *FLAWS-hco* was introduced to obtain a CSF-suppressed T1-weighted contrast with a low *B*1, *T*2* and *M*0 sensitivity. An additional study conducted at 7T showed that the *FLAWS-hco* contrast could be used in a similar manner as the *MP2RAGE-uni* contrast to measure T1 relaxation times [16]. However, to the best of our knowledge, the possibility of measuring T1 relaxation times from *FLAWS-hco* has never been validated for 3T imaging, thus preventing to combine both MP2RAGE and FLAWS imaging benefits in a single 3T sequence acquisition.

In this context, the current study aimed at 1) optimizing the FLAWS sequence at 3T using a Cartesian phyllotaxis k-space undersampling to decrease the sequence acquisition time; and 2) assessing, via *in-silico, in-vitro* and *in-vivo* experiments, the possibility of measuring the T1 relaxation times with the FLAWS sequence at 3T for both the linear and Cartesian phyllotaxis k-space undersampling acquisitions.

## 2. Materials and methods

### 2.1. Compressed sensing FLAWS sequence optimization

The aim of the FLAWS optimization performed in the current study was to obtain a FLAWS contrast close to the one previously proposed by Tanner et al. at 3T [3] with a 1 *mm* isotropic image resolution, a 256 × 240 × 192 *mm*^3^ field of view (FOV) and a 6 *mins* acquisition time. In this context, parallel imaging with a linear k-space undersampling could not be used, as a downsampling factor of 5 would be required to reach the sequence optimization aims –GRAPPA 5 reconstructions generated very noisy images in preliminary experiments (results not presented in this manuscript)–. The linear k-space undersampling was then replaced by a jittered Cartesian spiral phyllotaxis k-space undersampling that allowed for the reduction of the MP2RAGE acquisition time in a previous study [10]. The Cartesian phyllotaxis undersampling coordinates were determined using the following equation:

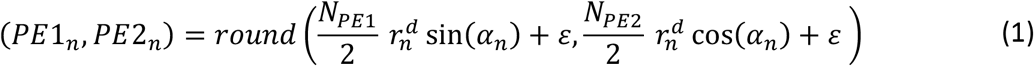

where:

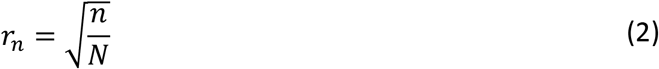

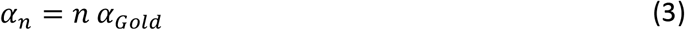

with *n* the current k-space sample, *N* the total number of k-space samples, *N*_*PE*1_ and *N*_*PE*2_ the number of phase encoding lines in the first and second phase encoding directions, *α_Gold_* the golden angle of the full circumference (approximately 137.5 °), *d* a density factor controlling the amount of samples near the k-space center and *ε* a random jitter used to add incoherence in the k-space sampling to facilitate the compressed sensing reconstruction [13]. Following the recommendations from Mussard et al. [10] to obtain accurate MP2RAGE reconstructions from the Cartesian phyllotaxis undersampling pattern, the density was set to 0.5 and the random jitter followed a Normal distribution with a mean of 0 and a variance of 1.2. The trajectories used for the k-space sampling were defined to ensure that the k-space center was sampled in the middle of the gradient echo train. The k-space sampling trajectories were determined as described in the Appendix section in [10] from *N*_*PE*2_, the number of samples per repetition time (TR) and the number of TR used to sample the k-space.

The FLAWS images were reconstructed from the undersampled k-space using a compressed sensing approach. First, coil sensitivity maps were computed with ESPIRIT [17] from fully sampled k-space data acquired with a FLASH sequence (acquisition time: 3 *secs*; matrix size: 32 × 32 × 64) [18]. As recommended by Mussard et al., the 3D image reconstruction problem was subdivided in multiple 2D image reconstruction problems to decrease the image reconstruction memory and computation time [10]. The reduction to 2D reconstruction problems was performed by computing the inverse Fourier transform along the frequency encoding direction. The 2D images were then reconstructed in the first and second phase encoding directions by minimizing the following equation:

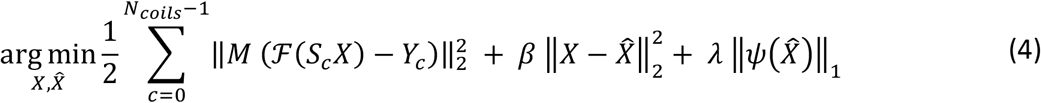

with *X* the image estimation, 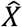 the image reconstructed after wavelet regularization, *N_colis_* the number of receiver coils, *M* the k-space sampling mask, 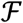 the discrete Fourier transform, *Y_c_* the acquired k-space data for a given coil *c, ψ* the wavelet transform and *β* and *λ* regularization parameters. In equation 4, the first L2 term is defined to ensure that the reconstructed image is consistent with the acquired data, the L1 term constrains the reconstructed image to be sparse in the wavelet domain and the second L2 term ensures that the images obtained from the L1 term and the first L2 term are consistent, thus allowing to split the reconstruction in two different subproblems [19]. Following the study conducted by Mussard et al. [10], the regularization parameter *β* was set to 0.4. The regularization parameter *λ* was manually tuned to obtain a good compromise between the noise reduction and the generation of blurring artefacts that arise according to the importance given to the sparsity of the reconstructed image in the wavelet domain.

The FLAWS1 and FLAWS2 images were reconstructed separately by minimizing equation 4. As the FLAWS1 and FLAWS2 images have different SNR, the regularization parameter *λ* was defined independently for both image reconstructions. In this manuscript, *λ*_1_ designs the regularization parameter *λ* used to reconstruct the FLAWS1 image and *λ*_2_ designs the one used to reconstruct the FLAWS2 image.

The FLAWS parameters were defined using a dedicated optimization method that was previously employed to optimize FLAWS at 1.5T and 7T [15,16]. This optimization method first consisted in selecting a set of pre-optimal parameters by maximizing a profit function under constraints that was devised to maximize the SNR in the FLAWS1 and FLAWS2 images, while ensuring a good CSF signal suppression in FLAWS2 and both a good WM signal suppression and visualization of deep GM structures in FLAWS1 [15]. Then, the optimal parameter set was selected among the pre-optimal parameter sets by maximizing the sum of the brain tissue contrasts in both FLAWS1 and FLAWS2 [15]. In line with previous studies, the globus pallidus was designed as the structure of reference for deep GM visualization in the profit function [15,16]. The profit function and contrast maximization steps of the optimization relied on FLAWS1 and FLAWS2 signal simulations based on the solutions of the Bloch equations corresponding to the FLAWS sequence. The WM and GM T1 relaxation times used for the signal simulations were the same as the ones used to optimize the FLAWS sequence at 3T in a previous study [3] (810 *ms* for WM and 1350 *ms* for GM). The globus pallidus T1 relaxation time was set to 901 *ms* [20] and the T1 relaxation time of the CSF was set to 4000 *ms*. Following previous FLAWS optimization studies, the proton densities were set to 0.7 for WM, 0.72 for the globus pallidus, 0.8 for GM and 1 for CSF [15,16]. An exhaustive search was used to define the optimal FLAWS sequence parameters, with the flip angles, *α*_1_ and *α*_2_, ranging from 4° to 13° (step-size: 1°); the first inversion time, *TI*_1_, ranging from the shortest first inversion time that can be obtained –the shortest first inversion time depends on the number of samples per TR and the gradient echo (GRE) repetition time– to 1500 *ms* (step-size:20 *ms*); and the second inversion time, *TI*_2_, ranging from the shortest second inversion time that can be obtained –the shortest second inversion time depends on the first inversion time, the number of samples per TR and the GRE repetition time– to the longest second inversion time (step-size: 20 *ms*) –the longest second inversion time depends on the sequence TR, the number of samples per TR and the GRE repetition time–. The GRE repetition time was set to 5.8 *ms* (the value of the GRE repetition time was constrained by the bandwidth used for the acquisition, which was set to 260 *Hz/Px*).

The number of samples per TR and the sequence TR affect both the k-space sampling percentage and the FLAWS1 and FLAWS2 contrasts. The FLAWS optimization was then performed multiple times with different sequence TR values (ranging from 3 *secs* to 5.5 *secs*, step-size: 0.5 *secs*) and number of samples per TR (100, 116 and 133). The choice of the number of samples per TR was constrained by the range of desired inversion times and acceleration factors (please refer to [10] for further details about the constrains on the number of samples per TR). The sequence TR and number of samples per TR were chosen to obtain the best trade-of between the percentage of the k-space that was sampled and the theoretical signal to noise ratio (SNR) and contrast loss compared to a reference FLAWS optimization.

The reference FLAWS optimization was obtained by using the FLAWS dedicated optimization method presented above for a linear k-space undersampling with a 6/8 partial Fourier, a fixed sequence TR of 5 *secs* and a fixed number of samples per TR of 144. Please note that these fixed parameters correspond to a standard FLAWS sequence optimization with a 1 *mm* isotropic resolution, a FOV of 256 × 240 × 192 *mm*^3^ and an acquisition time under 10 *mins* when using a linear k-space undersampling with a GRAPPA 3 reconstruction. In other words, the optimized parameters for the reference FLAWS fulfilled the compressed sensing optimization constrains except for the acquisition time which was longer than 6 *mins*.

#### 2.1.1. FLAWS contrasts and T1 mapping

The FLAWS contrasts and T1 maps were reconstructed as described below by using the *FLAWS-Tools* open-source software (*https://github.com/jerbeaumont/FLAWS-Tools*). The *FLAWS-min* (GM-specific), *FLAWS-mip* (GM-specific with low *B*1 sensitivity), *FLAWS-uni* (GM-suppressed with low *B*1 sensitivity), *FLAWS-hc* (WM-suppressed with low *B*1 sensitivity) and *FLAWS-hco* (CSF-suppressed with low *B*1 sensitivity) contrasts were reconstructed using the following equations:

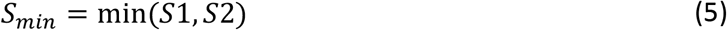

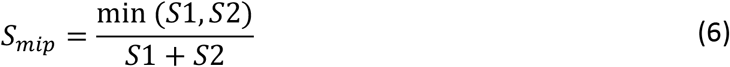

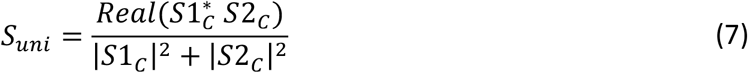

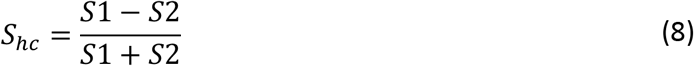

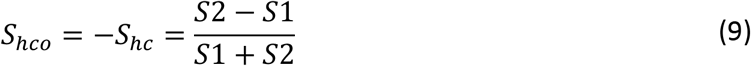

with *S*1 and *S*2 the magnitude of the *FLAWS1* and *FLAWS2* signals; *S*1_*C*_ and *S*2_*C*_ the complex *FLAWS1* and *FLAWS2* signals; and * the complex conjugate operator. T1 relaxation times were measured with the FLAWS sequence by building lookup tables of the *FLAWS-hc* signal as described in [16].

#### 2.1.2. In-silico, in-vitro and in-vivo MRI experiments

MR experiments were conducted on a 3T whole body MRI research scanner (Magnetom Prisma, Siemens Healthcare, Erlangen, Germany) equipped with a 64-channel head coil. *In-vivo* experiments were conducted on 10 healthy volunteers (27 to 46 years of age, 6 women). All experiments were performed under written informed consent and were approved by the institutional review board (Royal Brisbane and women’s hospital human research ethics committee, HREC/2020/QRBW/63943). Two different imaging protocols were used in the current study. The first protocol contained three different CS FLAWS optimizations –the CS optimizations were obtained from three different pairs of chosen sequence TR and number of samples per TR, designed as CS-P1, CS-P2 and CS-P3 in the manuscript– and the reference GRAPPA FLAWS optimization. In addition, the reference FLAWS parameters were used to obtain FLAWS images reconstructed from a full k-space sampling. The second protocol contained the CS-P1, CS-P2, GRAPPA and full k-space FLAWS sequences as well as an MP2RAGE sequence that was used as a reference for T1 mapping. In addition, a *B*1+ map was acquired in both protocols from a turbo fast low-angle-shot sequence (acquisition time: 32 *secs*; voxel size: 3.1 × 3.1 × 4 *mm*^3^; 3 measurement repeats) [21]. The parameters corresponding to the MP2RAGE and FLAWS sequences acquired in the two imaging protocols are presented in Table 1. Four healthy volunteers underwent imaging with the first protocol, four other volunteers were imaged with the second protocol and two volunteers were scanned with both protocols.

**Table 1.**
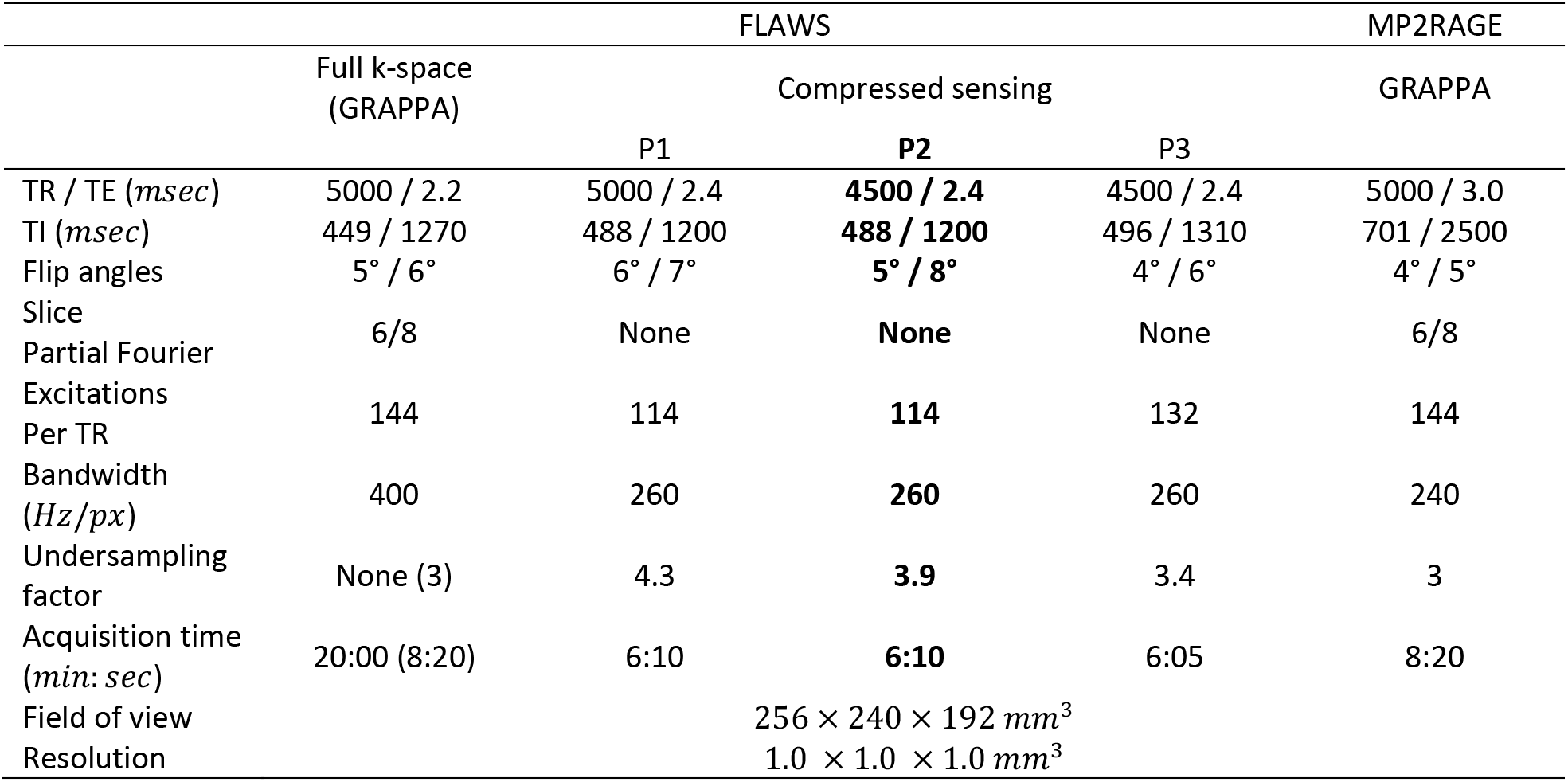
Sequence parameters used in the current study. The advised set of parameters for the FLAWS with compressed sensing acquisition is the one corresponding to the protocol P2 (bold).

The FLAWS optimizations were qualitatively validated *in-vivo* by assessing: the overall image quality; the quality of the WM and CSF signals suppression in FLAWS1 and FLAWS2, respectively; and the visualization of the separation between the internal and the external globus pallidus in FLAWS1. A quantitative *in-vivo* validation of the FLAWS optimizations was performed by measuring the contrast (CN) and the contrast to noise ratio (CNR) between brain tissues with the following equations:

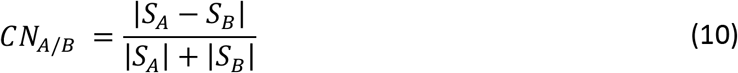

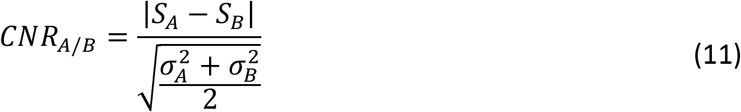

with *S_A_* (respectively *S_B_*) and *σ_A_* (respectively *σ_B_*) the mean and standard deviation of a given tissue *A* (respectively *B*). To be consistent with the results reported by Tanner et al. regarding the first optimization of the FLAWS sequence at 3T [3], the CN and CNR were measured between the WM, GM and CSF signals by manually drawing regions of interest in the corpus callosum (splenium), the caudate nucleus (head) and the lateral ventricles.

The theoretical accuracy and precision of the FLAWS and MP2RAGE T1 mappings were determined *in-silico* using Monte-Carlo experiments obtained from signals simulated as follows:

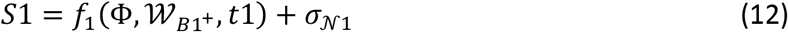

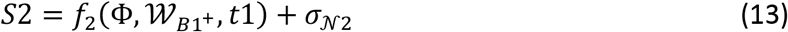

with *S*1 (respectively *S*2) the MP2RAGE1/FLAWS1 (respectively MP2RAGE2/FLAWS2) signal magnitude; *f*_1_ (respectively *f*_2_) the solution of the MP2RAGE/FLAWS sequence Bloch equations used to simulate MP2RAGE1/FLAWS1 (respectively MP2RAGE2/FLAWS2); Φ the set of MP2RAGE/FLAWS sequence parameters; 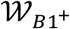 a random variable following a Weibull distribution (*α* = 9.36, *β* = 0.91, *μ* = −0.0029) tailored to fit the brain *B*1^+^ values measured in the current study (please refer to the Supplementary Materials from [16] for further information regarding the *B*1^+^ distribution fitting); *t*1 the T1 value used to simulate the MP2RAGE/FLAWS signals (T1 range: 700 – 2500 *ms*, step-size: 1 *ms*, 100 simulations per T1 value); and 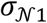 and 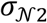 random variables following a normal distribution with a mean of 0 and a standard deviation determined to simulate a WM signal with a SNR of 25 in MP2RAGE2. The equations used to determine the theoretical T1 mapping accuracy and precision are provided in Supplementary Materials.

*In-vitro* experiments were conducted on an MRI caliber phantom (*QalibreMD* System Standard Model 130, https://qmri.com/system-phantom) containing 10 spheres with different T1 relaxation times. This phantom was scanned with the second imaging protocol on the 3T whole body MRI research scanner used for *in-vivo* experiments using a 20-channel head and neck matrix coil (the phantom did not fit into the 64-channel head coil used for the *in-vivo* experiments). In addition to the second imaging protocol, nine inversion recovery turbo spin echo (IRTSE) images were acquired with the following parameters: *TE* = 8.2 *ms*, TB = 6000 *ms*, *TIs* = 35/500/1000/1500/2000/2500/3000/4000/5000 *ms*, 1 *mm* isotropic resolution, total acquisition time: 52: 30 *mins*. T1 mapping was performed with all sequences. The MP2RAGE and FLAWS T1 maps were post-hoc corrected for *B*1^+^ inhomogeneites as described in [22]. The relative error between the FLAWS/MP2RAGE T1 maps and the IRTSE T1 map were computed in ROIs manually drawn in the 10 phantom spheres. T1 mapping was also performed on the *in-vivo* data acquired in this study. Similarly to the *in-vitro* experiments, the *in-vivo* T1 maps were post-hoc corrected for *B*1^+^ inhomogeneities [22]. The *in-vivo* T1 relaxation times were measured in the WM, putamen, caudate nucleus and cortical GM by manually drawing ROIs (the ROI drawing was performed as described in [23]). The MP2RAGE, GRAPPA FLAWS and CS FLAWS *in-vivo* T1 maps were spatially normalized onto the full k-space FLAWS T1 map using rigid registration of the second inversion images. The registrations were performed using the block-matching rigid registration algorithm implemented in the Anima software (Anima, RRID:SCR_017017; https://github.com/Inria-Visages/Anima-Public) [24,25]. The relative difference between the full k-space FLAWS T1 map and the other MP2RAGE and FLAWS T1 maps was computed to assess the presence of spatially varying T1 mapping errors that could occur between the different optimizations and reconstruction methods.

## 3. Results

### 3.1. Compressed sensing FLAWS sequence optimization

The set of full k-space and GRAPPA FLAWS parameters obtained with the optimization method described in section 2.1 are presented in Table 1. This set of parameters was used as a reference to compute the theoretical signal loss characterized by the CS FLAWS optimizations. The theoretical signal loss increased when the sequence TR decreased and when the number of samples per TR increased (Supplementary Table 1). Similarly, the sum of the brain tissue contrasts tended to decrease when the sequence TR decreased and the number of samples per TR increased. However, the k-space sampling percentage increased when the sequence TR decreased and the number of samples per TR increased. Three CS FLAWS optimizations that correspond to different pairs of sequence TR and number of samples per TR were selected to conduct MR experiments. These CS FLAWS optimizations are denoted as CP-P1, CS-P2 and CS-P3 FLAWS in the manuscript and are presented in Table 1. The CS-P1 FLAWS optimization was selected as it maximizes the k-space sampling percentage without leading to an average signal loss compared to the reference (full k-space/GRAPPA) FLAWS optimization (Supplementary Table 1). The CS-P2 FLAWS optimization increases the k-space sampling percentage (k-space sampling: 20.1 %) compared to the CS-P1 FLAWS (k-space sampling: 18.1 %), which leads to an average signal loss of 4 % compared to the reference FLAWS optimization. The CS-P2 FLAWS is also characterized by a small decrease in its sum of the brain tissue contrasts (*Σ_CN_* = 4.25) when compared to the CS-P1 FLAWS (*Σ_CN_* = 4.42). The CS-P3 FLAWS optimization further increases the k-space sampling percentage (k-space sampling: 23.1 %) to the cost of also further increasing the average signal loss compared to the reference FLAWS optimization (average signal loss: 14.3 %). The sum of the brain tissue contrasts obtained with the CS-P3 FLAWS optimization is close to the one obtained for the CS-P2 FLAWS (*Σ_CN_* = 4.24).

Visual inspection of the *in-vivo* CS FLAWS images reconstructed with *λ* parameters (range: 0.0001 – 0.0010, step-size: 0.0001) suggested to use *λ*_1_ = 0.0004 and *λ*_2_ = 0.0003 to obtain the best compromise between noise reduction and blurring artefact generation, for all CS FLAWS optimizations. These *λ* values were used to reconstruct the CS FLAWS images and associated results presented in this manuscript.

*In-vivo* experiments showed that all FLAWS protocols were characterized by a WM signal suppression in FLAWS1 and CSF signal suppression in FLAWS2, which validates the optimizations from a qualitative point of view (Figure 1). The three CS optimizations displayed a similar contrast to the one obtained with the full k-space and GRAPPA FLAWS optimizations (Figure 1). Visual inspection suggested that the full k-space FLAWS provided images with an overall better quality than the GRAPPA and CS FLAWS. In line with the theoretical signal loss computations, the visual inspection suggested that the CS-P3 FLAWS provided noisier images than the CS-P1 and CS-P2 FLAWS. No significant visual differences were noticed when comparing the GRAPPA FLAWS with the CS-P1, CS-P2 and CS-P3 FLAWS in terms of artefacts generated from the image reconstructions. The separation between the internal and the external globus pallidus was identified in FLAWS1 for all FLAWS protocols, thus further validating the protocols in a qualitative point of view. However, visual inspections suggested that the GRAPPA protocol provided a better globus pallidus visualization than the CS protocols (Supplementary Figure 1).

**Figure 1.**
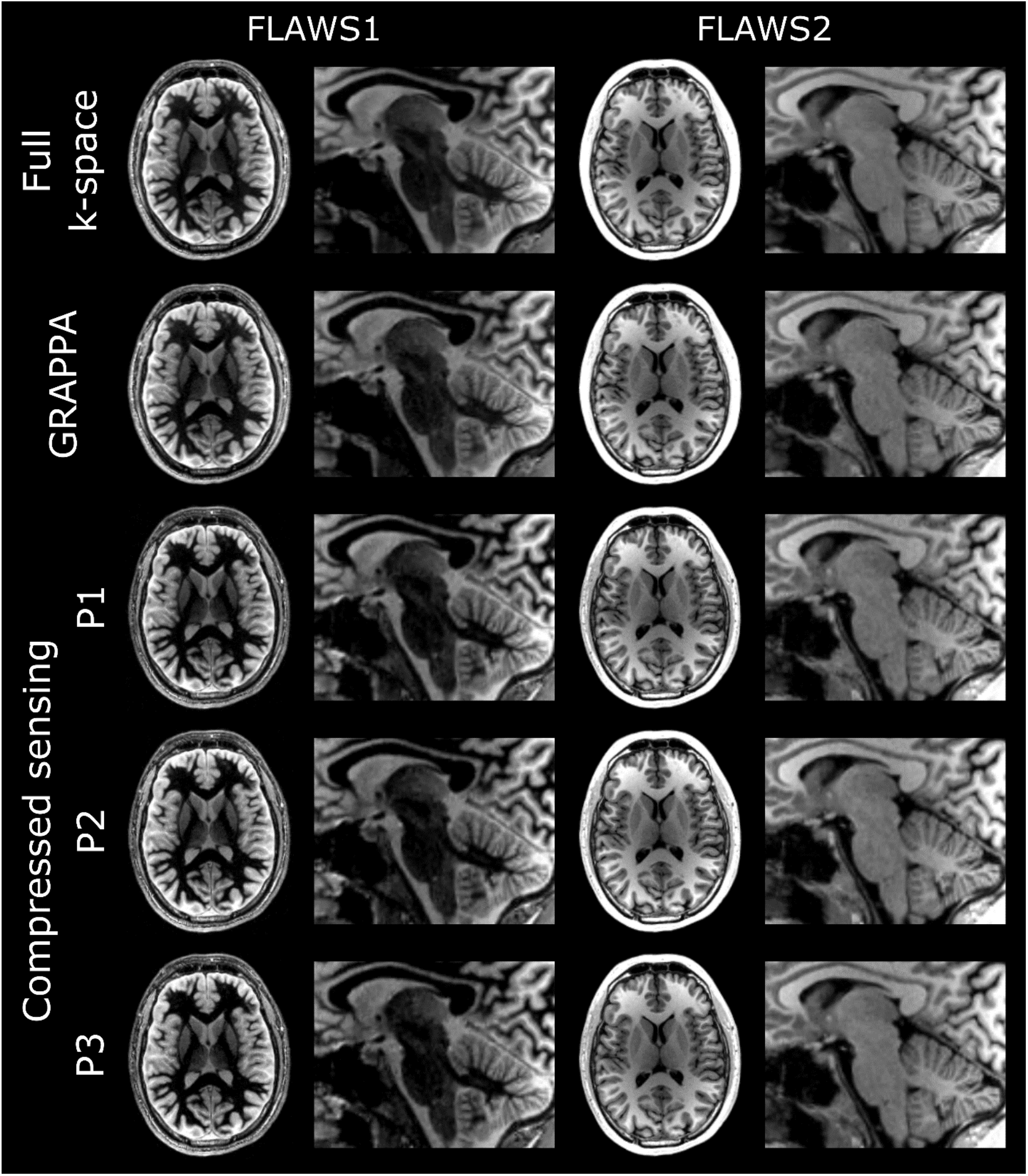
In-vivo FLAWS1 and FLAWS2 images obtained from the full k-space, GRAPPA and compressed sensing (CS, protocols P1, P2 and P3) FLAWS. All FLAWS provided a good white matter signal suppression in FLAWS1 and a good cerebrospinal fluid signal suppression in FLAWS2. The compressed sensing FLAWS displayed a brain tissue contrast that was close to the full k-space and GRAPPA FLAWS in both FLAWS1 and FLAWS2.

The seven FLAWS contrasts reconstructed from the CS optimizations (examples obtained for CS-P2 are shown in Figure 2) appeared visually close to the FLAWS contrasts obtained in previous studies conducted at 1.5T and 7T [14–16] and were also visually close to their counterparts that were reconstructed from the GRAPPA optimization. Specifically, a GM-specific contrast was obtained for *FLAWS-min* and *FLAWS-mip;* a WM-suppressed contrast was obtained for *FLAWS1* and *FLAWS-hc;* a CSF-suppressed contrast was obtained for *FLAWS2* and *FLAWS-hco;* and a GM-suppressed contrast was obtained for *FLAWS-uni*. In line with previous studies [15,16] and theoretical assumptions, the *FLAWS-hc, FLAWS-hco, FLAWS-mip* and *FLAWS-uni* contrasts were characterized by a low *B*1 sensitivity compared to *FLAWS1, FLAWS2* and *FLAWS-min (Figure 2*).

**Figure 2.**
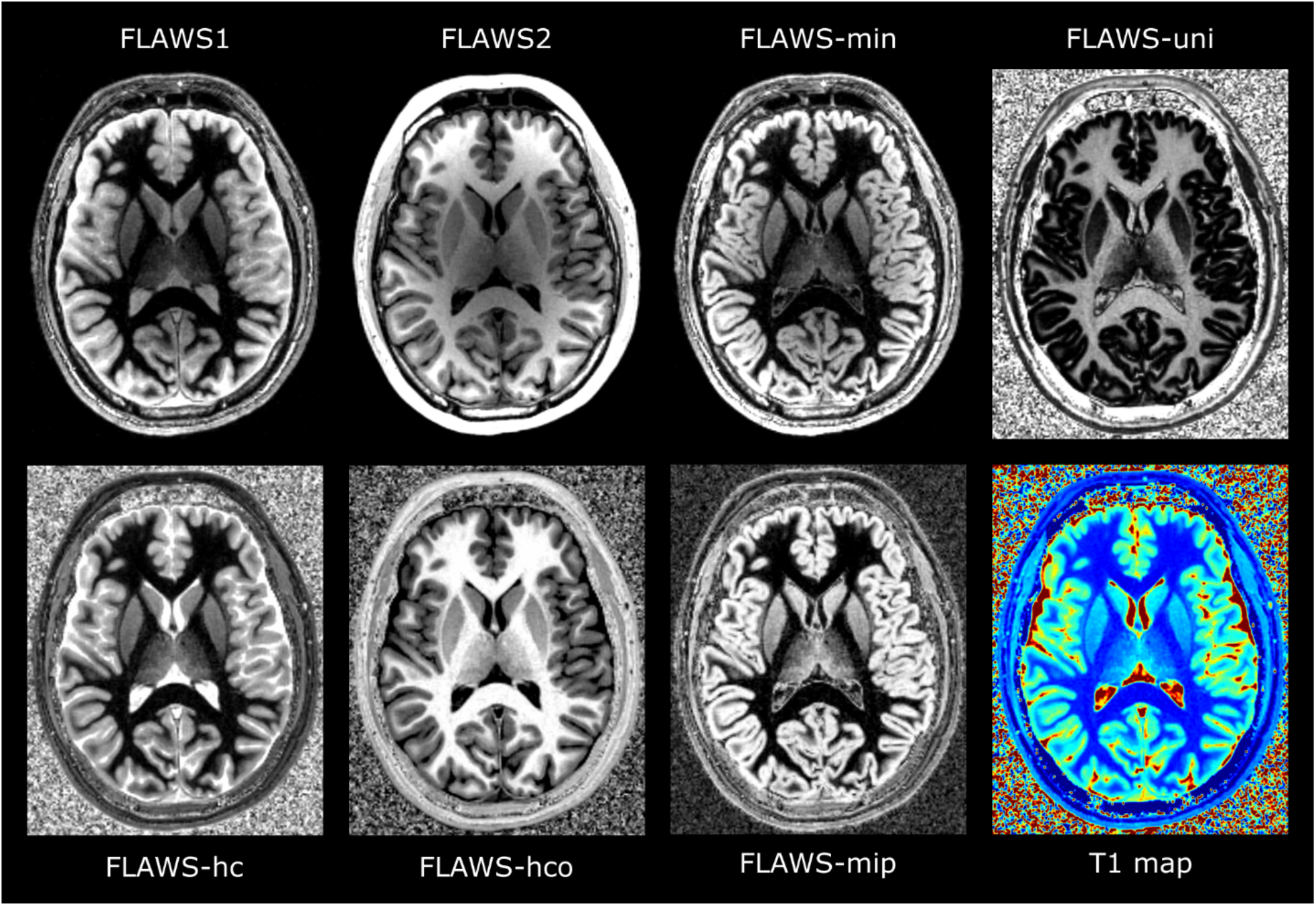
In-vivo FLAWS contrasts and T1 map obtained from the compressed sensing (CS) FLAWS corresponding to the protocol P2. The FLAWS1 and FLAWS-hc images displayed a brain tissue contrast characterized by a good white matter (WM) signal suppression. The FLAWS2 and FLAWS-hco contrasts provided a good cerebrospinal fluid (CSF) signal suppression. The FLAWS-min and FLAWS-mip images were characterized by a gray matter (GM) specific contrast. The FLAWS-uni image provided a GM-suppressed contrast. The FLAWS-hc, FLAWS-hco, FLAWS-mip, and FLAWS-uni images were characterized by a lower sensitivity to B1 field inhomogeneities compared to the FLAWS1, FLAWS2 and FLAWS-min images.

Quantitative measurements showed that the CS optimizations provided a contrast that was close to or higher than the GRAPPA optimization for all brain tissues (Table 2). In addition, both GRAPPA and CS optimizations provided a brain tissue contrast that was close to or higher than the one reported by Tanner et al. for the first FLAWS optimization conducted at 3T [3]. The CNR measurements provided results that were in line with the theoretical results and visual inspections: a higher CNR was measured for the full k-space FLAWS than for the GRAPPA and CS FLAWS (Table 3). The CS-P1 and CS-P2 FLAWS provided a CNR that was close to or higher than the GRAPPA FLAWS CNR. Please note that the higher CNR measured in the CS FLAWS compared to the GRAPPA FLAWS is partially due to the inherent image denoising performed during the CS image reconstruction. The CS-P3 FLAWS provided images with a lower CNR than the GRAPPA, CS-P1 and CS-P2 FLAWS. The CS-P3 optimization was discarded from further analysis as it was not considered as the best CS optimization due to its lower image quality from both qualitative and quantitative point of views. Since the CS-P1 and CS-P2 optimizations provided similar performances in terms of qualitative and quantitative image quality, the choice of the final CS optimization was determined according to its T1 mapping performances.

**Table 2.** In-vivo brain tissue contrasts measured in FLAWS1 and FLAWS2 for the GRAPPA FLAWS and the compressed sensing (CS) FLAWS acquisitions (protocols P1, P2 and P3). The brain tissue contrasts measured in the CS-P1, CS-P2 and CS-P3 FLAWS were close to or higher than the ones measured in the GRAPPA FLAWS for both FLAWS1 and FLAWS2. In addition, the FLAWS1 and FLAWS2 brain tissue contrasts obtained from the GRAPPA, CS-P1, CS-P2 and CS-P3 FLAWS optimizations were close to or higher than the ones obtained from the first FLAWS optimization conducted by Tanner et al. at 3T [3].

|  |  |  | WM/GM | WM/CSF | GM/CSF |
| --- | --- | --- | --- | --- | --- |
| FLAWS1 | GRAPPA | Tanner et al.<br>($n = 5$) | 0.59<br>(0.51 – 0.69) | 0.68<br>(0.62 – 0.77) | 0.16<br>(0.13 – 0.17) |
| | | This study<br>($n = 10$) | 0.70<br>(0.61 – 0.79) | 0.79<br>(0.70 – 0.85) | 0.19<br>(0.14 – 0.22) |
| | Compressed<br>sensing | P1<br>($n = 10$) | 0.75<br>(0.64 – 0.84) | 0.84<br>(0.77 – 0.89) | 0.25<br>(0.20 – 0.30) |
| | | P2<br>($n = 10$) | 0.74<br>(0.67 – 0.79) | 0.83<br>(0.78 – 0.85) | 0.21<br>(0.18 – 0.24) |
| | | P3<br>($n = 6$) | 0.71<br>(0.64 – 0.76) | 0.80<br>(0.75 – 0.83) | 0.19<br>(0.15 – 0.21) |
| FLAWS2 | GRAPPA | Tanner et al.<br>($n = 5$) | 0.15<br>(0.13 – 0.16) | 0.83<br>(0.68 – 0.89) | 0.78<br>(0.60 – 0.86) |
| | | This study<br>($n = 10$) | 0.16<br>(0.04 – 0.25) | 0.82<br>(0.76 – 0.85) | 0.76<br>(0.70 – 0.81) |
| | Compressed<br>sensing | P1<br>($n = 10$) | 0.17<br>(0.10 – 0.26) | 0.88<br>(0.84 – 0.93) | 0.84<br>(0.81 – 0.89) |
| | | P2<br>($n = 10$) | 0.17<br>(0.05 – 0.26) | 0.89<br>(0.82 – 0.93) | 0.85<br>(0.80 – 0.89) |
| | | P3<br>($n = 6$) | 0.15<br>(0.11 – 0.15) | 0.88<br>(0.84 – 0.90) | 0.84<br>(0.80 – 0.86) |

**Table 3.**
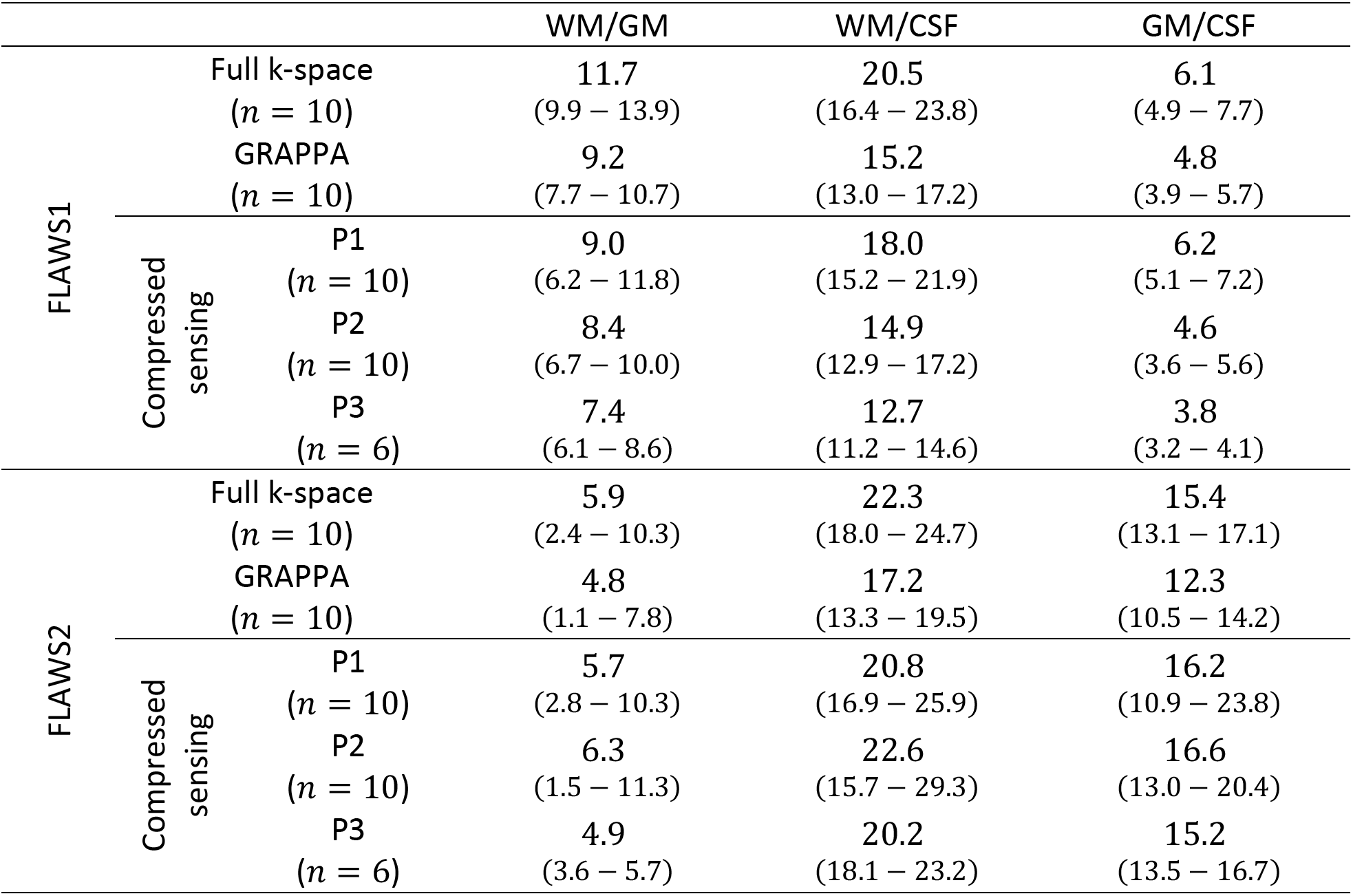
In-vivo brain tissue contrast to noise ratio (CNR) measured in FLAWS1 and FLAWS2 for the full k-space, GRAPPA and compressed sensing (CS, protocols P1, P2 and P3). The full k-space FLAWS was characterized by a higher CNR than the GRAPPA and CS FLAWS. The CS-P1 and CS-P2 FLAWS provided a higher brain tissue CNR than the GRAPPA FLAWS in both FLAWS1 and FLAWS2. The CS-P3 FLAWS provided a lower brain tissue CNR than the GRAPPA, CS-P1 and CS-P2 FLAWS.

#### 3.1.1. T1 mapping with the FLAWS sequence

Similarly to the MP2RAGE T1 mapping, the T1 mappings performed with the full k-space, GRAPPA and CS-P2 FLAWS were characterized by a low *B*1^+^ sensitivity in the 700 – 1800 *ms* T1 range (Supplementary Figure 2). The CS-P1 FLAWS was characterized by an increased T1 mapping *B*1^+^ sensitivity in this T1 range when compared to the full k-space, GRAPPA and CS-P2 FLAWS (Supplementary Figure 2). In line with a previous study conducted at 7T [16], the T1 mapping *B*1^+^ dependency obtained for the full k-space, GRAPPA, CS-P1 and CS-P2 FLAWS increased as a function of the T1 relaxation times, thus requiring a post-hoc *B*1^+^ correction to ensure accurate T1 measurements for long T1 relaxation times.

The raw T1 mapping accuracy and precision measured *in-silico* for the full k-space, GRAPPA and CS-P2 FLAWS were close to or higher than the MP2RAGE T1 mapping accuracy and precision (Table 4). The accuracy and precision obtained for the full k-space, GRAPPA and CS-P2 FLAWS were respectively higher than 95 % and 90 % in all simulation cases. The CS-P1 FLAWS provided a lower raw T1 mapping accuracy and precision than the full k-space, GRAPPA and CS-P2 FLAWS (Table 4).

**Table 4.** T1 mapping accuracy and precision measured in-silico for the MP2RAGE sequence and the full k-space, GRAPPA and compressed sensing (CS, protocols P1 and P2) FLAWS. The T1 mapping accuracy and precision obtained with the full k-space, GRAPPA FLAWS were close to the ones obtained with the MP2RAGE in every simulation cases. The CS-P2 FLAWS was characterized by performances close to the full k-space/GRAPPA FLAWS performances. The T1 mapping accuracy and precision obtained for the CS-P1 FLAWS were lower than the ones obtained for the MP2RAGE, full k-space, GRAPPA and CS-P2 FLAWS.

| Bias | SNR | Sequence | Accuracy (%) | Precision (%) |
| --- | --- | --- | --- | --- |
| No $B1^+$ | 25 | MP2RAGE | 99.6 | 90.4 |
|  |  | FLAWS full k-space/GRAPPA | 99.4 | 92.5 |
|  |  | FLAWS compressed sensing P1 | 99.4 | 92.9 |
|  |  | FLAWS compressed sensing P2 | 99.5 | 92.5 |
| $B1^+$ | $+\infty$ | MP2RAGE | 97.5 | 97.9 |
|  |  | FLAWS full k-space/GRAPPA | 95.4 | 96.2 |
|  |  | FLAWS compressed sensing P1 | 93.3 | 94.4 |
|  |  | FLAWS compressed sensing P2 | 95.3 | 96.1 |
|  | 25 | MP2RAGE | 97.1 | 88.7 |
|  |  | FLAWS full k-space/GRAPPA | 95.3 | 90.3 |
|  |  | FLAWS compressed sensing P1 | 92.9 | 90.1 |
|  |  | FLAWS compressed sensing P2 | 95.3 | 90.4 |

For all phantom spheres (T1 range measured with IRTSE: 287 – 2522 *ms*), a relative error below 5 % was found between the T1 relaxation times measured with IRTSE and the MP2RAGE, full k-space and GRAPPA FLAWS (Figure 3). The CS-P2 FLAWS provided T1 measurements with a relative error below 5 % for T1 relaxation times ranging from 771 *ms* to 2522 *ms* (Figure 3). However, the relative error between the CS-P2 FLAWS and IRTSE T1 measurements was higher than 5 % for T1 values lower than 771 *ms*. The CS-P1 FLAWS provided T1 measurements with a relative error that was higher than the CS-P2 FLAWS (Figure 3).

**Figure 3.**
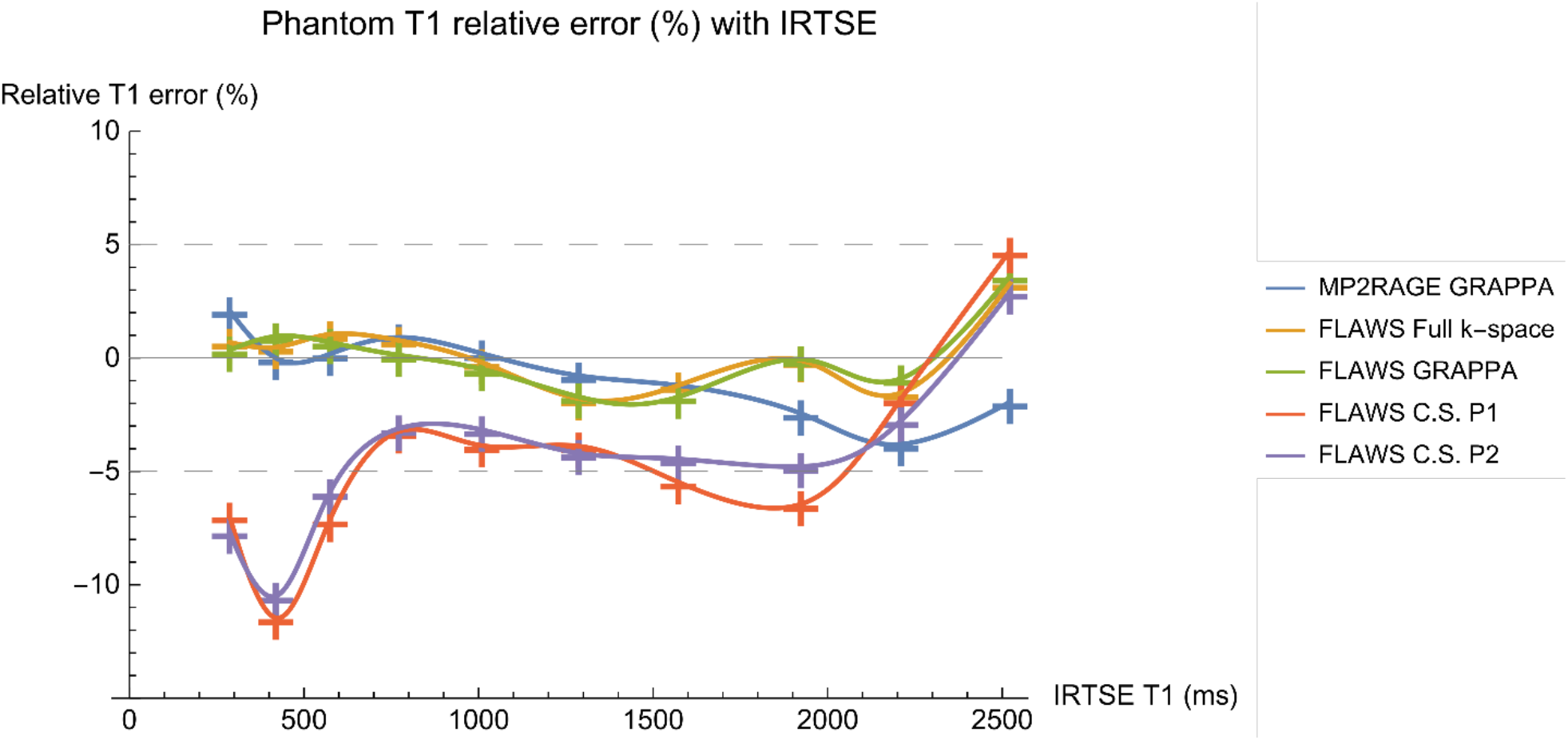
Relative error obtained in-vitro between the inversion recovery turbo spin echo (IRSTE) and the MP2RAGE/FLAWS T1 measurements. The relative T1 measurement error obtained from the full k-space and GRAPPA FLAWS was close to the one obtained from the MP2RAGE and was lower than 5% for all T1 measurements. The compressed sensing (CS, protocols P1 and P2) FLAWS provided a higher relative T1 measurement error compared to the full k-space and GRAPPA FLAWS. The relative T1 measurement error obtained from the CS-P2 FLAWS remained below 5% for T1 measurements between 700 ms and 2500 ms.

The relative error between the MP2RAGE T1 relaxation times and the full k-space, GRAPPA, CS-P1 and CS-P2 FLAWS T1 relaxation times measured *in-vivo* after post-hoc *B*1^+^ correction was close to or lower than 5 % in the WM, putamen, caudate nucleus and cortical GM (Table 5). The T1 relative difference in the WM and cortical GM ROIs was computed per subjects between the full k-space FLAWS and the sequences (Figure 4). The relative difference between the full k-space FLAWS T1 relaxation times and the MP2RAGE and GRAPPA FLAWS T1 relaxation times was below 5 % for all subjects. This difference was also below 5 % for most of the subjects for the CS-P1 and CS-P2 FLAWS (Figure 4). The relative T1 difference between the full k-space FLAWS and the other sequences was below 5 % for all subjects in the putamen and the caudate nucleus (Supplementary Figure 3). The T1 measurement error on repeated acquisitions was below 5 % for the full k-space, GRAPPA, CS-P1 and CS-P2 FLAWS in the WM, putamen, caudate nucleus and cortical GM on the two subjects that underwent both imaging protocols presented in section 2.3 (Figure 4, Supplementary Figure 3).

**Table 5.** In vivo T1 relaxation times measured in white matter (WM), putamen, caudate and cortical gray matter (GM) with the MP2RAGE, full k-space, GRAPPA and compressed sensing (CS, protocols P1 and P2) FLAWS. The full k-space, GRAPPA, CS-P1 and CS-P2 FLAWS provided T1 measurements with a relative error close to or lower than 5% when compared to the T1 measurements obtained from the MP2RAGE.

|  |  | WM | Putamen | Caudate | Cortical GM |
| --- | --- | --- | --- | --- | --- |
| MP2RAGE T1 (ms)<br>(n = 6) |  | 819<br>±38 | 1195<br>±58 | 1228<br>±40 | 1484<br>±27 |
| FLAWS full k-space<br>(n = 10) | T1 (ms) | 810<br>±44 | 1214<br>±67 | 1253<br>±55 | 1524<br>±42 |
|  | MP2RAGE<br>rel. error | 1.1 % | 1.6 % | 2.0 % | 2.7 % |
| FLAWS GRAPPA<br>(n = 10) | T1 (ms) | 812<br>±43 | 1219<br>±56 | 1257<br>±48 | 1510<br>±26 |
|  | MP2RAGE<br>rel. error | 0.9 % | 2.0 % | 2.4 % | 1.8 % |
| FLAWS CS P1<br>(n = 10) | T1 (ms) | 787<br>±46 | 1231<br>±62 | 1278<br>±51 | 1560<br>±35 |
|  | MP2RAGE<br>rel. error | 3.9 % | 3.0 % | 4.1 % | 5.1 % |
| FLAWS CS P2<br>(n = 10) | T1 (ms) | 780<br>±48 | 1230<br>±58 | 1279<br>±54 | 1549<br>±24 |
|  | MP2RAGE<br>rel. error | 4.8 % | 2.9 % | 4.2 % | 4.4 % |

**Figure 4.**
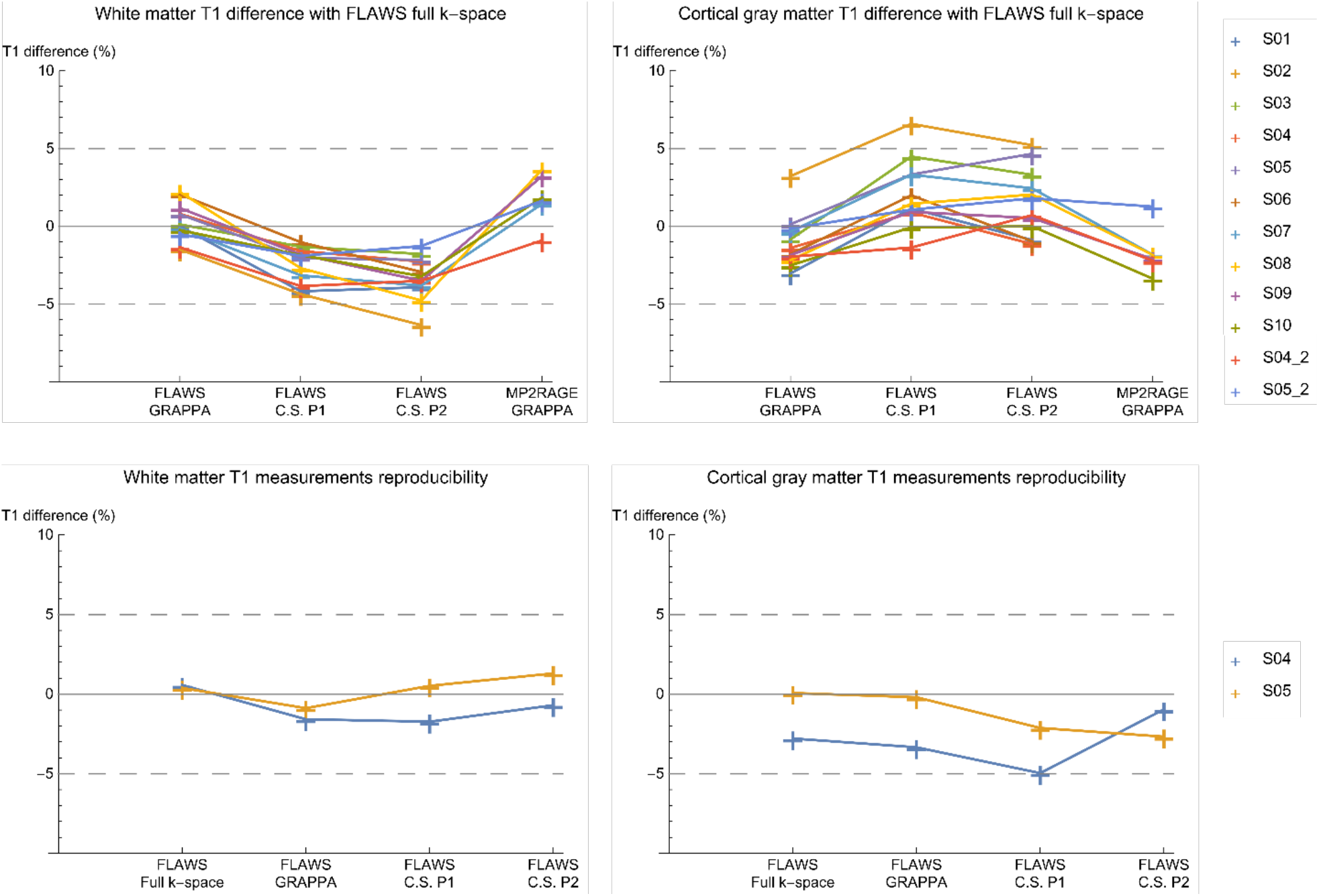
Relative T1 measurement difference obtained in-vivo for all subjects in white matter (top left) and cortical gray matter (top right) regions of interest (ROI) between the full k-space FLAWS and the MP2RAGE, GRAPPA and compressed sensing (CS, protocols P1 and P2) FLAWS. The relative T1 measurement difference between the full k-space FLAWS and the other acquisitions was close to or lower than 5% for most subjects. The T1 measurement error obtained on repeated acquisitions was lower than 5% for all acquisitions in the white matter (bottom left) and cortical gray matter (bottom right) ROIs.

The CS-P2 FLAWS provided T1 measurements that were closer to the reference T1s when compared to the CS-P1 FLAWS for in *in-silico* and *in-vitro* experiments. The difference obtained *in-vivo* between the CS-P2 FLAWS T1 and the reference T1 was close to or lower than the one obtained from the CS-P1 FLAWS. Therefore, the CS-P2 optimization was defined as the best CS optimization obtained in the current study as it provides the best results when considering both the overall FLAWS image quality and T1 mapping performances. Consequently, the CS-P1 FLAWS was discarded from further analysis in the current study.

A visual inspection suggested that the FLAWS T1 maps obtained after post-hoc *B*1^+^ correction appeared similar to their MP2RAGE counterparts (Figure 5). To assess the effect of the image reconstruction on T1 mapping, the relative difference between the full k-space FLAWS T1 map and the other T1 maps was computed after *B*1^+^ post-hoc correction (Figure 5). The highest differences between the T1 maps were found in the CSF and at the tissue borders, which were highlighted due to imperfect image co-registrations. The dominant difference between the full k-space FLAWS T1 map and the GRAPPA FLAWS and MP2RAGE T1 maps was below 5 %, while a higher dominant difference was found for CS-P2 FLAWS T1 map (close to 5 %). During visual inspection, the highest differences between the full k-space T1 maps and the CS-P2 FLAWS T1 maps did not appear to be linked to artefacts generated by the compressed sensing reconstruction, but where more linked to small co-registration errors between the T1 maps.

**Figure 5.**
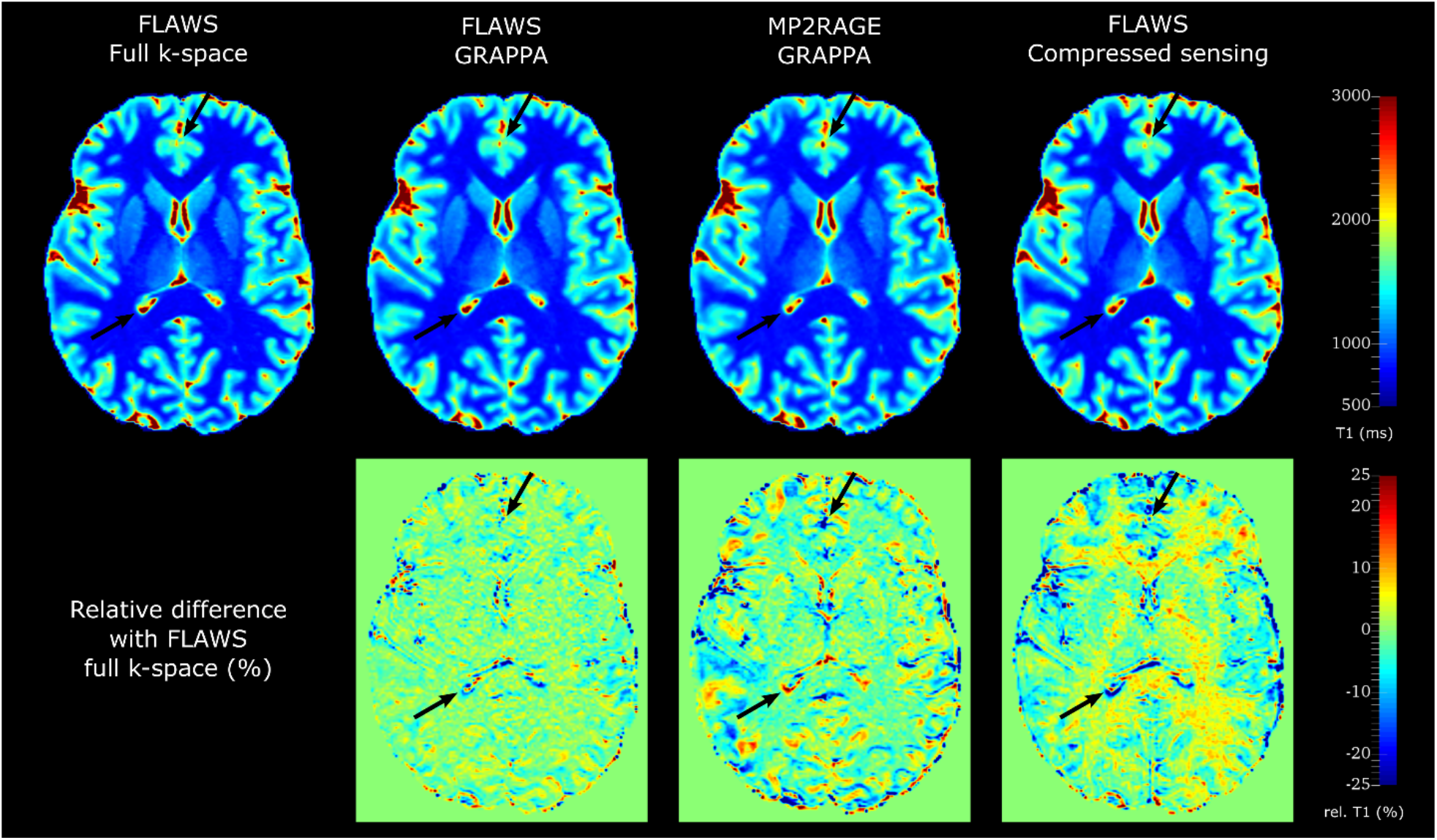
T1 maps obtained from the MP2RAGE, full k-space, GRAPPA and compressed sensing (CS, protocol P2) FLAWS. The T1 maps obtained from all acquisitions appeared visually similar to one another. The relative difference between the T1 map obtained from the full k-space FLAWS and the other T1 maps (bottom row) showed that the highest T1 measurement differences were located in the cerebrospinal fluid (CSF) or at the tissue borders. The high relative T1 differences obtained at the tissue borders (examples highlighted by the black arrows) were found to be due to imperfect image co-registrations. Overall, the MP2RAGE and GRAPPA FLAWS provided lower relative T1 differences with the full k-space FLAWS than the CS-P2 FLAWS.

## 4. Discussion

### 4.1. Compressed sensing FLAWS sequence optimization

The current study proposes an optimization of the FLAWS sequence at the field strength of 3T based on a cartesian phyllotaxis k-space sampling and a compressed sensing reconstruction. These k-space sampling and reconstruction strategies were used to decrease the FLAWS sequence acquisition time, which was prohibitive for clinical application in the case where linear k-space undersampling and GRAPPA reconstruction strategies were used. The choice of the k-space undersampling pattern was motivated by the fact that it successfully allowed to reduce the MP2RAGE sequence acquisition time in a previous study conducted at 3T [10]. Since the FLAWS sequence is an MP2RAGE pulse sequence with a different parameter optimization, the choice of the density and jitter parameters used in the cartesian phyllotaxis k-space undersampling pattern followed the recommendations given in the previous MP2RAGE study [10]. The remaining parameters were determined using a method based on a profit function maximization with constraints on the FOV, brain tissue contrast and sequence acquisition time.

The optimization method used in this study was selected as it allowed to successfully optimize the FLAWS sequence in previous studies conducted at 1.5T and 7T [15,16]. Three CS FLAWS parameter sets with different k-space sampling percentage and brain tissue CNR were obtained from the optimization. The choice of the best set of CS parameters was based on in-silico, in-vitro and in-vivo assessments regarding: 1) the brain tissue CNR in FLAWS1 and FLAWS2, 2) the generation of artefacts in the CS image reconstruction and 3) the T1 mapping performances.

According to the aforementioned criterions, the CS-P2 optimization is the best CS optimization proposed in this study. The CS-P2 FLAWS images were qualitatively and quantitatively similar to the GRAPPA FLAWS images and did not contain reconstruction artefacts that could have hampered clinical interpretation or quantitative analysis. The CS-P2 FLAWS provided an *in-vivo* contrast that was in line with the first FLAWS *in-vivo* contrast obtained by Tanner et al. at 3T [3]. Please note that the FLAWS sequence parameters proposed by Tanner et al. were not used as a reference for GRAPPA FLAWS imaging as they did not fulfill the FOV constraints of the current study.

One of the characteristics of FLAWS imaging is the visualization of the globus pallidus [3,15,16]. The CS-P2 FLAWS allows for the identification of the separation between the internal and the external globus pallidus, which was used as a qualitative validation metric in previous FLAWS optimization studies [3,15,16]. However, the CS-P2 FLAWS is characterized by a lower globus pallidus visualization quality compared to the GRAPPA FLAWS. This lower globus pallidus visualization quality can be explained by the longer inversion time obtained in the CS-P2 optimization, which is mainly constrained by the k-space sampling pattern and the number of samples per TR.

### 4.2. T1 mapping with the FLAWS sequence

T1 mapping was performed with the FLAWS sequence at 3T using a method previously proposed for 7T FLAWS T1 mapping [16]. To the best of our knowledge, the current study is the first study showing that T1 mapping can be performed with the FLAWS sequence at 3T. Specifically, the *in-silico, in-vitro* and *in-vivo* experiments performed in this study show a good agreement between the GRAPPA FLAWS T1 mapping and T1 mappings obtained from methods of reference in the 700 – 2500 *ms* T1 range, which covers the WM, deep GM and cortical GM tissues.

*In-vitro* and *in-vivo* qualitative and quantitative comparisons between the full k-space FLAWS and the GRAPPA FLAWS and MP2RAGE T1 mappings show a good agreement despite differences in their k-space sampling patterns. Specifically, the highest T1 mapping differences obtained appear to be mainly due to imperfect image co-registrations and do not seem to be caused by artefacts that could have been generated during the image reconstructions. Despite the fact that the CS-P2 FLAWS T1 mapping is characterized by slightly higher T1 mapping differences against reference measurements than the GRAPPA FLAWS, its performances are sufficient to allow for its use in *in-vivo* studies. Specifically, the CS-P2 FLAWS provides 1) a high *in-silico* T1 mapping accuracy and precision and 2) *in-vitro* and *in-vivo* measurements that agree with the reference measurements in the T1 range covering the WM, deep GM and cortical GM tissues. In agreement with the GRAPPA FLAWS, the CS-P2 FLAWS also provides spatial T1 mapping differences with the full k-space FLAWS that appear to be mainly due to imperfect image co-registration and that do not seem to be caused by the presence of wavelet artefacts that could have been generated during the CS reconstruction.

### 4.3. Limitations

This study has multiple limitations. First, only a small number of healthy volunteers underwent *in-vivo* imaging and more experiments should be performed on a large patient cohort to fully assess the potential of this work for inclusion in clinical practice. In addition, the choice of the parameters used for the CS reconstruction was either empirical or based on recommendations from a previous study conducted on CS MP2RAGE imaging. However, the chosen parameters allowed for the reconstruction of CS FLAWS images that were of similar quality to the GRAPPA FLAWS images and did not seem to contain wavelet artefacts that could have hampered clinical interpretation or quantitative analysis. Specifically, the *λ*_1_ and *λ*_2_ parameters were manually tuned in the CS reconstruction to obtain images with the best trade-off between denoising and wavelet artefact generation, with a specific care taken to minimize the presence of wavelet artefacts in the images generated.

The notion of image quality as presented in this study is subjective and no clear gold standard exists regarding the assessment of image quality in FLAWS image reconstruction. In the current study, the image quality was determined through qualitative and quantitative assessments that were in line with previous studies reported in the literature [3,10,15,16]. In the previous study providing a CS optimization of the MP2RAGE sequence [10], Mussard et al. investigated the volume changes between the GRAPPA and CS MP2RAGE acquisitions in different brain structures to further assess image quality. A limitation of the current study is that volumetric changes were not taken into account in our image quality assessments. The absence of volumetric measurements is explained by the fact that, to the best of our knowledge, the gold standard brain parcellations techniques such as Freesurfer [26] have not yet been validated for FLAWS imaging. Therefore, using such methods to assess volumetric changes would first require to validate their use on FLAWS data, which would exceed the scope of this work. Volumetric assessments should however be carried in future studies to further validate the use of Cartesian phyllotaxis k-space sampling and CS reconstruction for FLAWS imaging at 3T.

## 5. Conclusion

In conclusion, this study proposes a FLAWS sequence optimization tailored to allow for the acquisition of FLAWS images with a Cartesian phyllotaxis k-space sampling and CS reconstruction. It was shown that this new CS FLAWS optimization allows to reduce the FLAWS sequence acquisition time from 8 *mins* to 6 *mins* without decreasing the FLAWS images quality. In addition, this study shows that T1 mapping can be performed with the FLAWS sequence for both GRAPPA and CS acquisitions. These new results suggest that the recent advances in FLAWS imaging [15,16] allow to combine both the MP2RAGE and the previous version of FLAWS imaging benefits in a single sequence acquisition.

## 6. Acknowledgements

We thank the “Region Bretagne” (France), which partially funded the current study. We are also grateful to the team at the Herston Imaging Research Facility for their generous scan time allowances and technical support acquiring the data presented here.

## Supplementary materials

### *In-silico* T1 mapping accuracy and precision

The *in-silico* T1 mapping accuracy and precision were computed as follow:

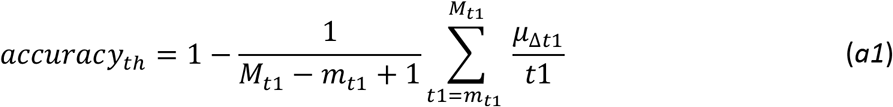

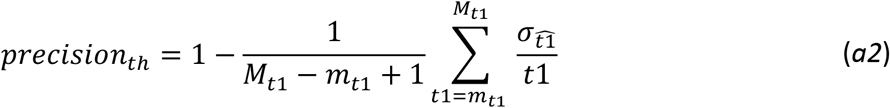

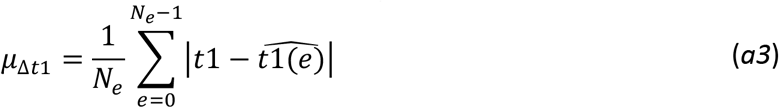

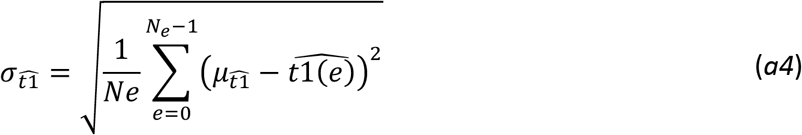

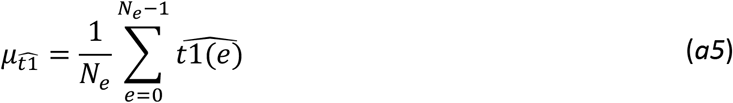

With *m*_*t*1_ (resp. *M*_*t*1_) the minimum (resp. maximum) T1 value used to generate the Monte-Carlo experiments, *N_e_* the number of Monte-Carlo experiments, *t*1 the T1 value used to simulate a given Monte-Carlo experiment *e* and 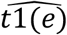 the T1 value estimated from a given Monte-Carlo experiment *e*.

**Supplementary Table 1.**
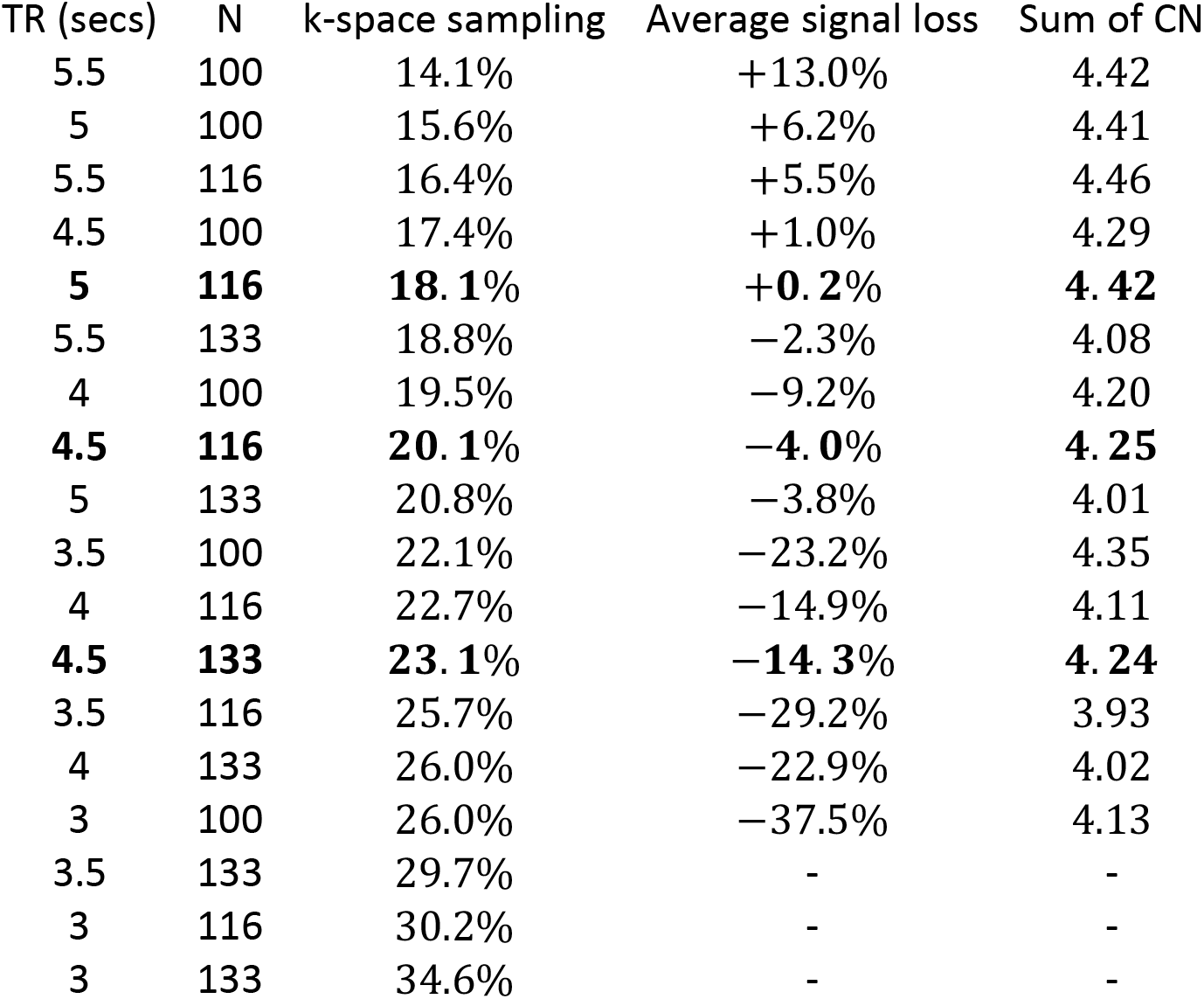
K-space sampling percentage, average signal loss compared to the reference FLAWS optimization and sum of brain tissue contrasts of the compressed sensing (CS) FLAWS optimizations obtained with different sequence repetition times (TR) and number of samples per TR (N). The k-space sampling percentage increases by reducing the sequence TR and increasing the number of samples per TR, while the average signal loss and sum of the brain tissue contrasts tend to decrease when the sequence TR decreases and the number of samples per TR increases. Three different CS FLAWS optimizations were selected for MR experiments (shown in bold in the table). CS-P1 corresponds to the set of parameters for TR = 5 secs and N = 116. This set of parameters prevents signal loss compared to the GRAPPA FLAWS optimization, to the cost of obtaining a lower k-space sampling percentage. CS-P2 corresponds to the set of parameters for TR = 4.5 secs and N = 116. This set of parameters is characterized by a small increase in the average signal loss, a decrease of the sum of the brain tissue contrast and a higher k-space sampling percentage than CS-P1. CS-P3 corresponds to the set of parameters for TR = 4.5 secs and N = 133. This set of parameters allows to further increase the k-space sampling percentage compared to CS-P2, to the cost of an increased average signal loss. The dashes design TR and N pairs for which no set of parameters fulfilled the constraints in the profit function maximization (see section 2.1 of the main manuscript).

**Supplementary Figure 1.**
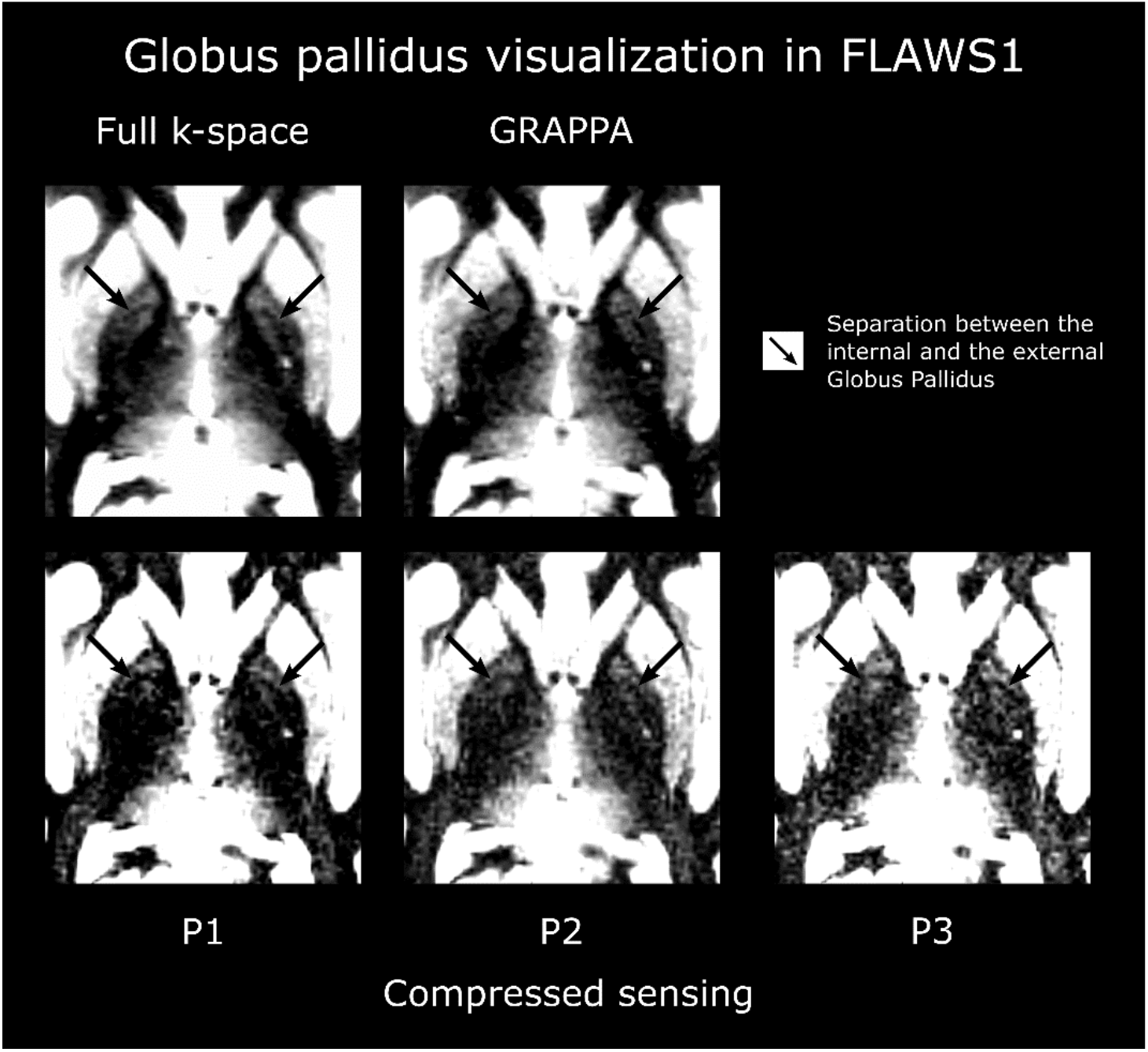
Globus pallidus visualization in FLAWS1 for the full k-space, GRAPPA and compressed sensing (CS, protocols P1, P2 and P3) FLAWS. The separation between the internal and the external globus pallidus was clearly identified for all FLAWS. However, the full k-space and GRAPPA FLAWS displayed a better globus pallidus visualization than the CS FLAWS.

**Supplementary Figure 2.**
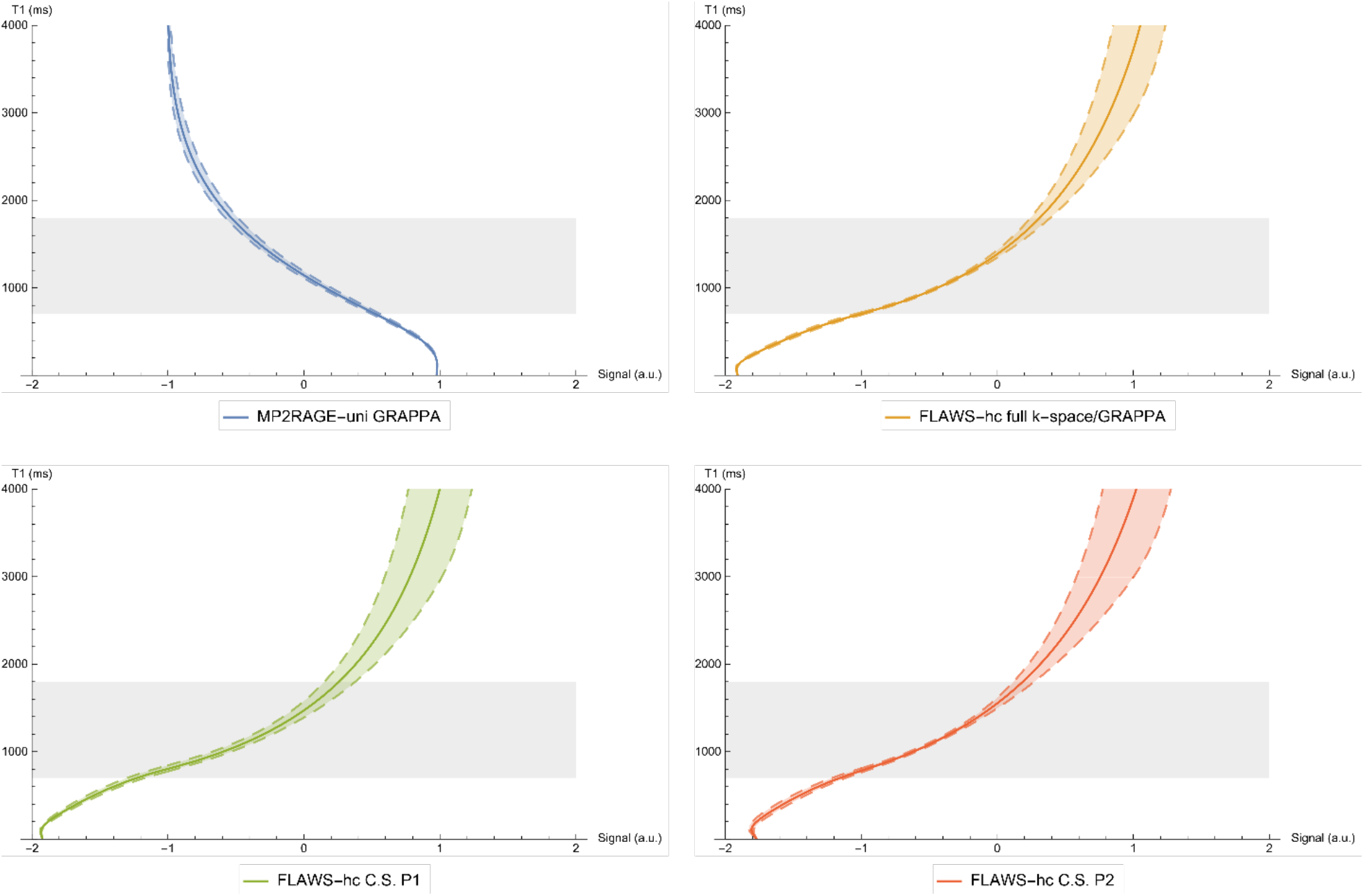
T1 relaxation times as a function of the MP2RAGE-uni and FLAWS-hc signals for the full k-space, GRAPPA and compressed sensing (CS, protocols P1 and P2) FLAWS. The dashed lines show the signals in the presence of ±20% B1^+^ inhomogeneities. The gray zone designs the 700 – 1800 ms T1 range, which covers the white matter, deep gray matter and cortical gray matter tissues. In agreement with the MP2RAGE, the full k-space, GRAPPA and CS-P2 FLAWS provided T1 relaxation time measurements with a low B1^+^ sensitivity in the 700 – 1800 ms T1 range. The B1^+^ sensitivity of the T1 mapping from the CS-P1 FLAWS was characterized by a higher B1^+^ sensitivity than the CS-P2 FLAWS. All FLAWS T1 mappings were characterized by an increased B1^+^ dependency as a function of the T1 relaxation times.

**Supplementary Figure 3.**
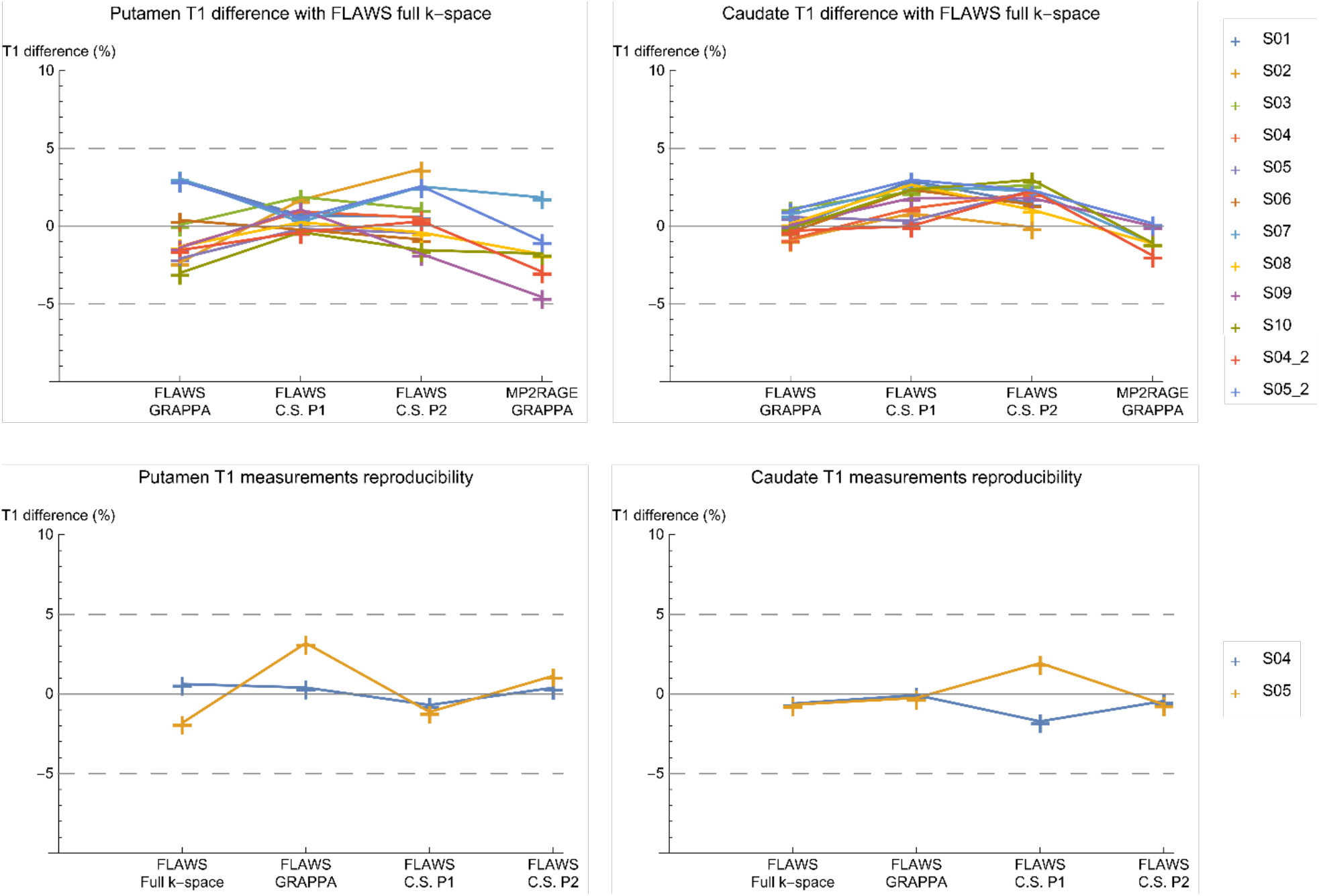
Relative T1 measurement difference obtained in-vivo for all subjects in the putamen (top left) and caudate (top right) regions of interest (ROI) between the full k-space FLAWS and the MP2RAGE, GRAPPA and compressed sensing (CS, protocols P1 and P2) FLAWS. The relative T1 measurement difference between the full k-space FLAWS and the other acquisitions was lower than 5 % for all subjects. The T1 measurement error obtained on repeated acquisitions was lower than 5 % for all acquisitions in the putamen (bottom left) and caudate (bottom right) ROIs.

